# A Deep Neural Network Two-part Model and Feature Importance Test for Semi-continuous Data

**DOI:** 10.1101/2023.06.07.544106

**Authors:** Baiming Zou, Xinlei Mi, James G. Xenakis, Di Wu, Jianhua Hu, Fei Zou

## Abstract

Semi-continuous data frequently arise in clinical practice. For example, while many surgical patients suffer from varying degrees of acute postoperative pain (POP) post surgery (i.e., POP score *>* 0), others experience none (i.e., POP score = 0), indicating the existence of two distinct data processes at play. Existing parametric or semi-parametric two-part modeling methods for this type of semicontinuous data can fail to appropriately model these two underlying data processes as such methods rely heavily on (generalized) linear additive assumptions. However, many factors may interact to jointly influence the experience of POP non-additively and non-linearly. Motivated by this challenge and inspired by the flexibility of deep neural networks (DNN) to accurately approximate complex functions universally, we derive a DNN-based two-part model by adapting the conventional DNN methods by adding two additional components: a bootstrapping procedure along with a filtering algorithm to boost the stability of the conventional DNN, an approach we denote as sDNN. To improve the interpretability and transparency of sDNN, we further derive a feature importance testing procedure to identify important features contributing to the outcome measurements of the two data processes, denoting this approach fsDNN. We show that fsDNN not only offers a valid feature importance test but also that using the identified features can further improve the predictive performance of sDNN. The proposed sDNN- and fsDNN-based twopart models are applied to the analysis of real data from a POP study, in which application they clearly demonstrate advantages over the existing parametric and semi-parametric two-part models. Further, we conduct extensive numerical studies to demonstrate that sDNN and fsDNN consistently outperform the existing two-part models regardless of the data complexity. An R package implementing the proposed methods has been developed and deposited on GitHub (https://github.com/SkadiEye/fsDNN).

## 1. Introduction

In many clinical settings, outcomes are often semi-continuous measurements, taking a boundary value (e.g., zero) under one condition, or a (positive) value arising from a continuous (or ordinal) distribution under another. This ubiquitous semicontinuous type of data tends also to be highly skewed with a spike at zero (Taylor, 1995; Farewell et al., 2017; Sauzet et al., 2019). For example, while many surgical patients suffer from acute postoperative pain (POP), defined as pain experienced within one month post surgery, with varying degrees of severity (POP intensity *>* 0), other patients experience no such pain (POP intensity = 0) (Gan, 2017). Such semi-continuous data abound in many other clinical and biomedical settings. For example, a subset of mice develop cancer after infection with Listeria monocytogenes, while others never do. Correspondingly, the tumor mass for the cancer-free mice is measured at exactly zero, while the value for those who do contract cancer will be a positive number (Boyartchuk et al., 2001). Examples of such data are manifold in economic studies for example, in studies of medical expenditures and length of hospital stays. In a given year some people accrue no medical expense (or require no time in hospital) at all. Otherwise, these outcomes are represented by positive numbers (i.e., dollars or days). Further, the size of an insurance claim will be a positive number when an incident occurs, and zero otherwise. Epidemiology studies provide more examples still. For example, many preterm-born children incur adverse health effects and are indexed by positive health outcome scores to indicate the corresponding adverse health outcome severity, while many other preterm-born children do not experience any adverse health effects and their corresponding health outcome index scores are recorded as zeros (Bangma et al., 2018, 2019).

Over 60% of the 100 million patients worldwide who have surgery each year suffer from moderate to severe acute POP (Kehlet, Jensen and Woolf, 2006). Despite great strides made by the scientific and clinical communities to understand POP mechanisms and treatments, inadequately controlled acute POP remains a major healthcare problem, associated with functional impairment, delayed recovery time, and prolonged duration of opioid use (Lovich-Sapola, Smith and Brandt, 2015). Poorly managed acute POP can also lead to extended length of hospital stay, increased risk of hospital readmission, and the development of chronic POP, defined as pain experienced by patients more than three month after surgery (Kehlet, Jensen and Woolf, 2006; Chou et al., 2016; Nair et al., 2020). Precise prediction of acute POP status and severity is crucial to help physicians apply timely and targeted pain treatments and to develop effective pain management plans to achieve earlier mobilization (Nair et al., 2020; Bayman, Oleson and Rabbitts, 2021). It guides clinicians toward the safest and most effective preemptive and preventative analgesic interventions, permitting the application of perioperative analgesic therapies in a manner that optimizes the benefit to risk ratio (Tighe et al., 2015a). However, existing acute POP forecasting methods do not lead to appropriate clinical decisions due to their failure to capture the complexity of acute POP, which restricts their predictive abilities (Tighe et al., 2015a).

A two-part mixture model is routinely employed to analyze the type of semi-continuous data that pertains to POP. In the existing model framework, logistic regression is frequently adopted to model the binary events (i.e., zero versus positive values), while a parametric (e.g., multiple linear or Poisson regression) (Duan et al., 1983; Min and Agresti, 2002; Mihaylova et al., 2011; Belotti et al., 2015) or semi-parametric method (Zhou and Liang, 2006) is used to model the non-zero portion of the data. In these modeling strategies, the observed clinical and demographic factors and additional biomarkers are often modeled in (generalized) linear additive fashion. However, this restrictive assumption often does not hold and is in any case difficult to verify in practice (Harrell, Lee and Mark, 1996; Beck and Jackman, 1998; Hastie, Tibshirani and Friedman, 2001; Che et al., 2019). Moreover, existing POP studies have suggested that many factors such as age, gender, anesthesia method, and surgery type and duration all interact to affect the likelihood of experiencing acute POP and its subsequent intensity (Trikha and Singh, 2013; Tighe et al., 2015b, 2016; Yang et al., 2019). Furthermore, these factors often influence the experience of acute POP in intricate ways, such as through the combination of their linear, quadratic, and even more complicated interaction terms (Zheng et al., 2017; Bayman, Oleson and Rabbitts, 2021).

The complex etiology of acute POP is difficult to model using existing parametric and semi-parametric two-part models, as these require explicit specification of the association between risk factors and disease outcomes. Typically, (generalized) linear additive assumptions are employed to model these associations, which are not flexible enough to capture complex effects like interactions and nonlinearities, which can lead to poor prediction accuracy. Motivated by this challenging problem and inspired by the flexibility of deep neural network (DNN) models for universally and accurately approximating complex functions, we herein derive a DNN-based two-part predictive model for semi-continuous data. The appeal of this technique is that it enables the robust modeling of highly complex functional relationships via layer-by-layer convolutions of outputs, using a subjects’ baseline profiles at the input layer to achieve very accurate predictions (Hornik, Stinchcombe and White, 1989; Farago’ and Lugosi, 1993; Tu, 1996; Sonoda and Murata, 2017). The flexibility and accuracy of DNN models to approximate complex functions is ensured by the universal approximation theorem (Cybenko, 1989; Hornik, 1991; Lu et al., 2017), although they can be unstable in finite sample settings, a challenge we address by incorporating bootstrap aggregating (bagging) and filtering in a method we denote sDNN. Bagging has been shown to boost the performance of unstable procedures, like those used in regression trees and neural networks (Hansen and Salamon, 1990; Breiman, 1996; Kotsiantis, 2014). However, due to the random parameter initialization in conventional DNN, there is no guarantee that each bagged dataset will necessarily lead to a stable optimal model in the finite sample setting. In neural networks, a guiding maxim is that “many could be better than all” (Zhou, Wu and Tang, 2002), which implies that using only top-performing bootstrapped DNN models could potentially improve the final ensemble model (Zhou, Wu and Tang, 2002; Mi, Zou and Zhu, 2019). Based on this argument, we further derive a scoring algorithm to filter out those poorly performing DNN models, leading to a final ensemble model that includes only those with good predictive performance.

Though sDNN-based two-part models can flexibly model the complex functional relationship and greatly improve prediction accuracy for semi-continuous outcomes, it is difficult to interpret each factor’s role in the two data processes. To overcome this disadvantage, we further develop a permutation-based feature importance testing procedure to correctly identify important features associated with the semi-continuous outcome, which we denote fsDNN. Permutation-based feature importance testing procedures have been shown to be powerful tools for identifying important features for different data types using various machine learning models (Breiman, 2001; Altmann et al., 2010; Putin et al., 2016; Mi et al., 2021). Using the identified features can further improve the prediction accuracy of our sDNN-based two-part model. We apply the sDNN- and fsDNN-based two-part models to a POP dataset that motivated this research, and compare them with the existing parametric and semi-parametric two-part predictive models.

The remainder of the paper is arranged as follows. In Section 2, we provide a detailed description of the proposed sDNN modeling procedures that adapt the conventional DNN models to include the bootstrapping and filtering algorithm. Additionally, we describe the details of fsDNN, our permutation-based testing procedure for feature importance identification. In Section 3, we then apply the sDNN and fsDNN-based two-part models to predict the acute POP measurements in our motivating dataset to demonstrate their benefits over the existing parametric and semi-parametric methods. Further, we conduct extensive numerical studies in Section 4, drawing comparisons with the conventional methods under different data complexity settings to explore the comparative applicability and performance of our methods. We demonstrate the generalizability of our sDNN and fsDNN-based twopart predictive models to a very general class of semi-continuous data. The paper concludes with some discussion in Section 5.

## 2. DNN-based Two-part Predictive Model and Feature Importance Test

Before presenting the details of the proposed sDNN and fsDNN framework, we introduce the models and notation. Let **X** represent all the observed clinical, demographic and biomarker data, and *Y* represent the outcome of interest, which can take a zero or a positive (e.g., continuous, ordinal) value. Our primary scientific interest is to model Pr(*Y >* 0 | **X**) and E (*Y* |*Y >* 0, **X**) flexibly and robustly such that we can obtain precise predictions for future patients. That is, we seek to predict both the propensity of a future patient to develop the disease, and the ensuing disease severity based on an observed baseline profile **X**.

We can express E(*Y* |**X**) = Pr(*Y >* 0|**X**)E (*Y* |*Y >* 0, **X**). For *Y* |{*Y >* 0, **X**}, we model E (*Y* |*Y >* 0, **X** = **x**) ≡ *ξ*(**x**). Further, we introduce *Z* = **I**(*Y >* 0) with **I** being an indicator function, and define Pr(*Z* = 1|**X** = **x**) ≡ *π*(**x**). Traditionally, a two-part parametric regression framework is routinely used to model this type of zero-inflated data (Duan et al., 1983; Min and Agresti, 2002; Mihaylova et al., 2011; Belotti et al., 2015). That is, *π*(**x**) and *ξ*(**x**) are assumed to be functions of linear combinations of all observed covariates **X** and estimated via, for example, logistic regression and multiple linear or Poisson regression models, respectively. To relax the parametric distributional assumption, Zhou and Liang (2006) extended the parametric modeling of *ξ*(**x**) for the positive portion of the data using a semi-parametric method employing a single-index model with an unspecified link function. That is, *ξ*(**x**) is assumed to be an unspecified function of linear combinations of the observed covariates **X**. However, the (generalized) linear additive assumptions are not flexible enough to capture the potential complex interactions and nonlinear effects for practical acute POP data. Herein, we relax these restrictive assumptions with unspecified functional forms for *π*(**x**) and *ξ*(**x**), and instead generate estimates of them (denoted as 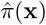 and 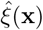, respectively) via the DNN approach presented in the following subsection. Based on these estimates, we predict the acute POP propensity and severity for a future patient with basel ine profile **X**_*new*_ = **x**_*new*_ as: 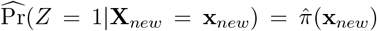 and 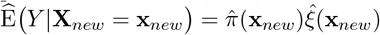, respectively.

### 2.1. Scoring Algorithm for Constructing Stable DNN Ensemble

To allow robust modeling of acute POP outcomes (i.e., POP status of zero vs. positive, and POP severity when POP status is positive) given a patient’s observed baseline profile **X** = **x**, i.e., Pr(*Z* = 1|**X** = **x**) ≡ *π*(**x**) and E (*Y* |*Y >* 0, **X** = **x**) ≡ *ξ*(**x**), we first adapt an *L*-hidden-layer feedforward DNN model (LeCun, Bengio and Hinton, 2015) to approximate the unspecified functions *π*(**x**) and *ξ*(**x**). For each hidden layer *l* ∈ {1, … *L*}, the model takes the input from the previous layer (which we denote as ***h***^(*l*−1)^, a vector with *n*_(*l*−1)_ elements), and transforms it through the function ***g***_*l*_(.) to generate the output ***h***. That is, ***h*** = ***g***_*l*_ ***b*** + ***W h*** ^−^, where ***b***^(*l*)^ is the bias vector of length *n*_*l*_ and ***W*** ^(*l*)^ is an *n*_*l*_ ×*n*_*l*−1_ weight matrix. The function ***g***_*l*_ can be regarded as applying the activation function *g*_*l*_ element-wise to the *n*_*l*_ dimensional vector ***b***^(*l*)^ + ***W*** ^(*l*)^***h***^(*l*−1)^. This activation function is a non-linear function that transforms the output values of neurons in the previous layer into the input values of the next layer. We take the common approach using a single function *g* for all the *g*_*l*_’s (*l* = 1, …, *L*); specifically, we employ a rectified linear unit (ReLU) function (Cybenko, 1989). For the first hidden layer (i.e., *l* = 1), ***h***^(0)^ is simply the original *p*-dimensional feature matrix ***X***. Thus, ***g***_1_ takes a *p*-dimensional input and produces an *n*_1_-dimensional output. Finally, the *L*^th^ hidden layer ***h***^(*L*)^ is connected to the output *Y* (*Z*) through *Y* (*Z*) ∼ *g*_out_ ***b***^(*L*+1)^ + ***W*** ^(*L*+1)^***h***^(*L*)^, where *g*_out_ gives a scalar output. The final output function is as the follows:

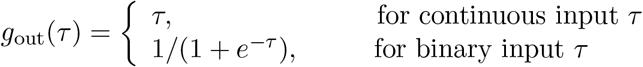

Thus, the final convolution output is *f* = *g*_out_ ◦ ***g***_*L*_ ◦ · · · ◦ ***g***_1_, where *f* takes the profile ***X*** as input and contains 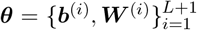 as a collection of parameters. Under this setup, ***θ*** are estimated by minimizing the following empirical risk function,

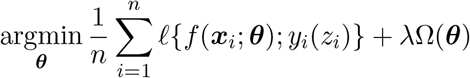

where *f*(·; ·) is the loss function, Ω(***θ***) is a penalty function and *λ* is a hyper-parameter that controls the degree of regularization. For the continuous outcome *Y* = *y* | *y >* 0, we set the loss function *f*(*f, y*) to 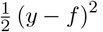. For a binary event *Z* = *z, f*(*f, z*) is set to the cross-entropy loss, − {*z* log *f* + (1 − *z*) log (1 − *f*)}. Furthermore, to shrink the model size, we put an *l*_1_ regularizer on the weight matrices ***W*** (*l* = 1, …, *L* + 1), i.e., 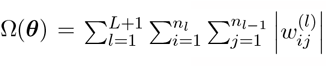, where 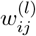 is the *ij*^th^ element in the weight matrix ***W*** ^(*l*)^. We optimize the risk function by using the mini-batch stochastic gradient descent algorithm (Byrd et al., 2012; Mei, 2018), along with an adaptive learning rate adjustment method called adaptive moment estimation (Adam) (Kinga and Adam, 2015) to obtain the parameter estimate 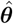 and the corresponding prediction at the profile 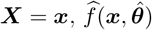.

### Bootstrap Aggregating Procedure

For a DNN model, the total number of parameters in ***θ*** is 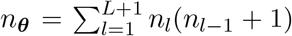, which is usually greater than the sample size *n*, leading to over-parameterization and unstable prediction. The accuracy of the prediction from a single DNN model (i.e., 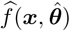) is therefore expected to be unreliable when the sample size is finite. An ensemble of neural networks has been shown to outperform a single neural network in such scenarios (Hansen and Salamon, 1990). Thus, we adopt bootstrap aggregating, i.e., bagging (Breiman, 1996), to increase the stability and accuracy of the DNNs.

We generate bootstrap samples by randomly sampling the training set with replacement *K* times. For each sample, we fit a DNN model, reserving the unused, a.k.a., the out-of-bag (OOB) samples as a validation set. Minimization of the risk function is performed for each bootstrapped sample to obtain parameter estimates, which are denoted as 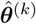 for the *k*^*th*^ sample. Further, 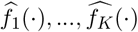 correspond to the fitted models for each of the *K* bootstrap samples. To aggregate the predictions from the *K* fitted models, a natural choice of the aggregated bagging prediction for a new observation with feature profile input ***X*** = ***x*** is given as 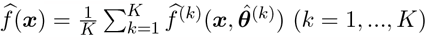.

### Filtering Procedure

Application of the bagging technique has been shown to substantially improve the robustness and accuracy of DNN predictions. However, due to the random parameter initializations used to fit these models, some DNNs may not converge to a stable solution and thus perform poorly. In neural network ensembles, it is argued that “many could be better than all” (Zhou, Wu and Tang, 2002), which suggests that using a subset of bagged DNNs that fit the data well could be better than using all of them (Zhou, Wu and Tang, 2002; Mi, Zou and Zhu, 2019). Therefore, we propose to remove those poorly performing DNNs from the final ensemble model, according to the criteria defined below. For the *k*^th^ bootstrap sample, a performance score *v*_*k*_ is defined as

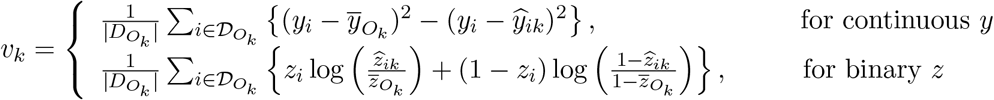

where 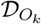 is the set of OOB samples, 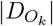 is the associated number of OOB samples, 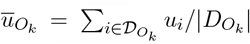 and 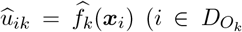, and *u* = *y* or *z*). For the regression DNN, *v*_*k*_ is the mean squared error loss, while for the classification DNN, *v*_*k*_ can be interpreted as the negated binomial deviance.

To determine the optimal subset of DNNs to be retained in the final ensemble, we first rank the DNNs based on their performance scores, denoting *v*_(1)_ ≥ · · · ≥ *v*_(*K*)_. Aggregating the top *q* DNNs, the prediction for ***x***_*i*_ (*i* = 1, · · ·, *n*) is then

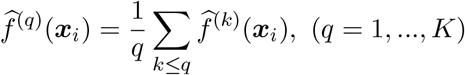

where 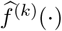 is the fitted DNN corresponding to the performance score *v*_(*k*)_. The optimal number of DNNs utilized by the ensemble, *q*_opt_, is determined by minimizing the training loss such that

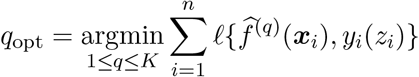

based upon which we obtain the revised bagging prediction 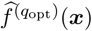 for a new observation with input profile ***X*** = ***x***.

In summary, in the proposed scoring ensemble algorithm, DNNs are ranked based on their performance in the OOB samples, and the optimal number of DNN models is determined by minimizing the loss in the training samples. The scoring algorithm not only can improve the stability of the DNN method, but also further reduces the potential for model overfitting. We refer to this revision of the DNN ensemble method as sDNN (for *stable* DNN), and summarize it in the following algorithm:

#### Algorithm 1

sDNN: Stable DNN for Two-part Predictive Modeling

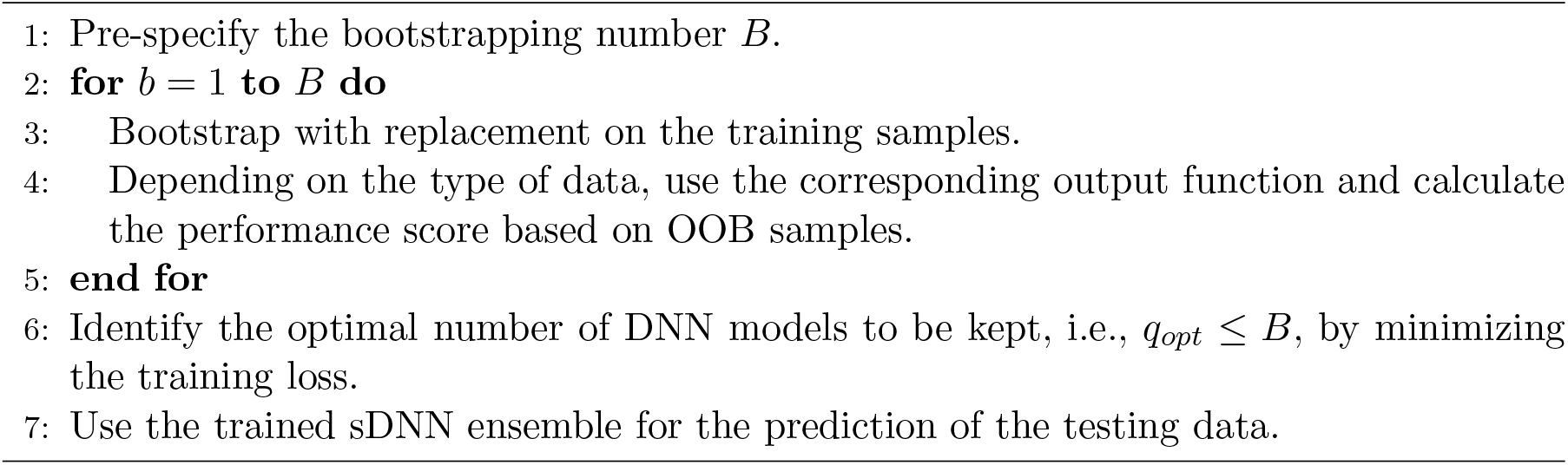

### 2.2. Feature Importance Test for sDNN Two-part Model

To determine important features contributing to the positive data process among all the p-dimensional features **X** under *E Y Y >* 0, **X** = **x** *ξ*(**x**), we define the feature importance score *M*_*j*_ of *X*_*j*_ (i.e., the *j*^th^ feature in **X**, *j* = 1, …, *p*) as the expected squared difference between *ξ*(**X**) and *ξ* **X**^(*j*)^, where **X**^(*j*)^ = (*X*_1_, …, *X*_*j*−1_, *X*_*j*_, *X*_*j*+1_, …, *X*_*p*_), i.e., **X** with its *j*^*th*^ feature replaced by a random permutation of the elements of *X*_*j*_. The importance score *M*_*j*_ can be rewritten as 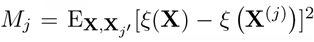, which is zero only when *ξ*(**X**) = *ξ*(**X**^(*j*)^), implying no contribution of **X**^(*j*)^ on *ξ*(**X**) conditional on the other features. The larger the impact of **X**^(*j*)^ on *ξ*(**X**), the larger *M*_*j*_ is expected to be. Also, *M*_*j*_ can be further decomposed as *M*_*j*_ = E_**X**,**X**_ [*Y* − *ξ*(**X**^(*j*)^)]^2^ − E_**X**_[*Y* − *ξ*(**X**)]^2^ and be estimated empirically. Let 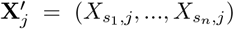 be a random sample of the elements in *X*_*j*_ without replacement, and the empirical permutation importance score be 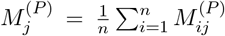 where 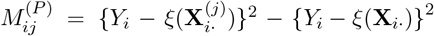 with **X**_*i*·_ = (*X*_*i*1_, …, *X*_*ip*_) and 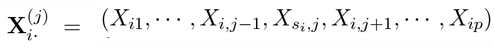. Note that 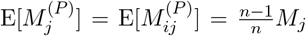. Letting 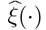 be the estimation of *ξ*(·) using sDNN method as described above, we propose to estimate 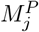 as 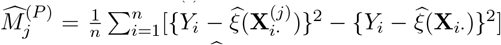. To avoid a potential overfitting of the sDNN approximation 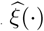 under finite sample size, we employ a cross-fitting strategy to separate the input data into training and validation sets, with the training set used for generating 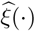 and the validation set for estimating 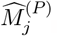. Let 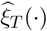 be the estimate of *ξ*(·) from the training set, and 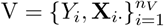 be the validation set; we have

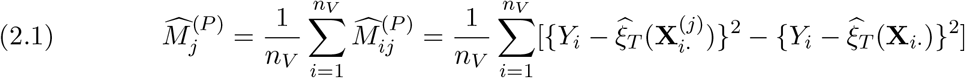

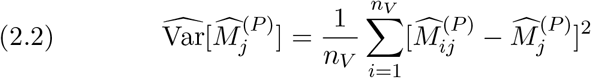

Based on it, we can construct test statistics as 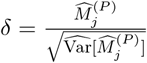, to determine each feature’s role in predicting the positive outcome *Y* and derive the important features.

To identify the important features for binary outcome *Z* with *π*(**X**) = E(*Z*|**X**) = Pr(*Z* =1|**X**), we defin e the im portance scor e *M*_*j*_ as th e expectation of binomial deviance, 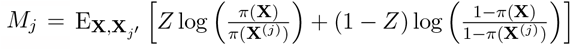. The empirical estimate of 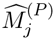, can be similarly obtained by plugging in the estimate of *π*(**X**^(*j*)^) and *π*(**X**) using sDNN method with cross-fitting strategy as described in continuous outcome scenario. Similarly, we can obtain the empirical estimate of Var 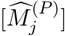, i.e., 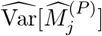, as given below:

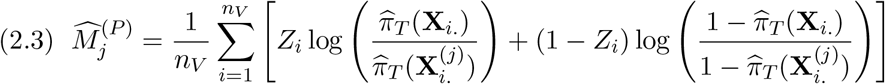

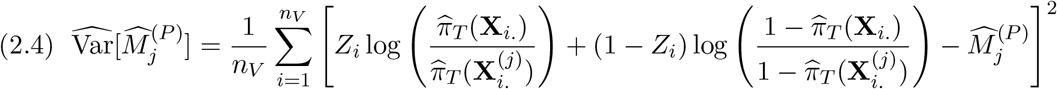

Test statistics 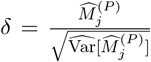 can be constructed to obtain the set of important features associated with binary event Z. Steps to identify important features for semi-continuous outcomes under sDNN two-part predictive modeling framework are summarized in Algorithm 2.

## 3. Application of sDNN and fsDNN Two-part Modeling to Acute POP Data

We apply the proposed sDNN and fsDNN two-part predictive modeling framework as described above to the clinical data from the POP study that motivated this research (Tighe et al., 2016). In this study, a clinical interest was to classify patients’ acute POP status and to predict the maximum pain intensity experienced 72 hours (i.e., on the 4^*th*^ day) postsurgery. This study had a total of 7, 391 patients and collected patient profile information on 37 variables, including anesthesia method, surgical duration and type, age, gender and body mass index (BMI). The distributions of some of the important variables are summarized in Table 1. On the fourth day 4, 135 (55.9%) patients achieved absence of acute POP, while 3, 256 patients still suffered from acute POP of varying degrees of intensity.

**Table 1:** Distribution of Major Clinical and Demographic Variables for Surgery Patients

| Categorical Variable |  | N | Proportion |
| --- | --- | --- | --- |
| Gender | Female | 3743 | 0.506 |
|  | Male | 3648 | 0.494 |
| Ethnicity | Hispanic | 206 | 0.028 |
|  | Non-Hispanic | 6890 | 0.932 |
|  | Unknown | 295 | 0.040 |
| Surgery Type | Cardiovascular | 965 | 0.131 |
|  | Digestive | 1556 | 0.211 |
|  | Integumentary | 519 | 0.070 |
|  | Musculoskeletal | 1746 | 0.236 |
|  | Nervous | 1043 | 0.141 |
|  | Pulmonary | 273 | 0.037 |
|  | Urinary | 569 | 0.077 |
|  | Other | 720 | 0.097 |
| Anesthesia Method | Nerve Block | 4004 | 0.542 |
|  | Partial Nerve Block | 1443 | 0.195 |
|  | General Anesthesia | 1944 | 0.263 |
| Opioid Use | Yes | 2381 | 0.322 |
|  | No | 3741 | 0.506 |
|  | Unknown | 1269 | 0.172 |
| Continuous Variable |  | Mean | SD |
| Age |  | 56.04 | 16.15 |
| BMI |  | 29.13 | 7.24 |
| Surgery Duration |  | 3.64 | 2.07 |

In addition to sDNN and fsDNN, we applied two other existing methods to the data. The first is a parametric method (referred to as Parametric), in which logistic regression is used to model the binary event of acute POP occurrence, and multiple linear regression is employed to model the continuous positive portion of the data, with all observed covariates entered as linear terms in these models (Duan et al., 1983; Olsen and Schafer, 2001; Min and Agresti, 2002; Mihaylova et al., 2011). The second existing method is a semi-parametric approach (referred to as Semiparametric) (Zhou and Liang, 2006) wherein a logistic regression model is used for the binary events (with all observed covariates entered as linear terms as in the Parametric approach) and a single-index model with unspecified link function is used for modeling the positive continuous portion data including the linear combination of all the observed baseline covariates.

### Algorithm 2

fsDNN: Permutation Feature Importance Test for sDNN Two-part Model

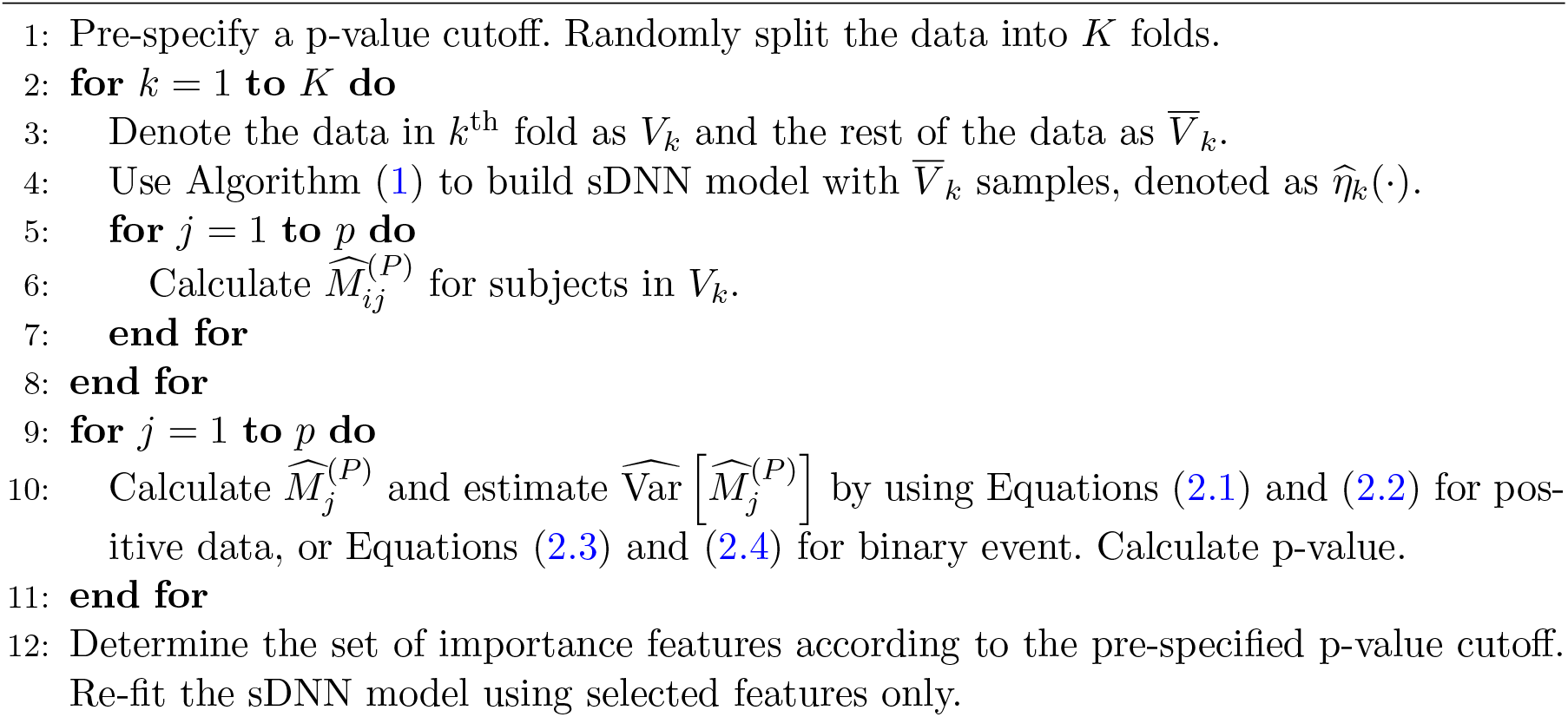

We evaluated all the methods considered based on their classification accuracy for (binary) pain status and prediction accuracy for (continuous) POP intensity via 10-fold crossvalidation. For classification, we define Accuracy 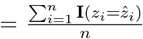, where 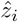 is the predicted binary outcome for sample *i* and **I** is the indicator function. The prediction of the binary outcome is given as follows:

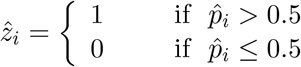

where 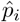 is the predicted probability that sample *i* experiences positive acute POP, based on the corresponding method employed. In addition to this metric, we use the area under the receiver operating characteristic curve (AUC) metric to compare classification accuracy. To evaluate the continuous outcome predictions we use the mean squared prediction error (MSPE) for both the conditional and marginal expectations of the testing samples. For the conditional expectation, we define 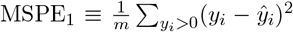 where *ŷ*_*i*_ is the predicted continuous outcome for sample *i*, and *m* is the number of samples with positive *y*. For the marginal expectation, we use 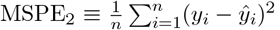. For all methods considered, negative predictions were replaced with zeros. All models included the 37 collected baseline clinical and demographic variables. For the proposed sDNN and fsDNN methods, we used neural network structure (30, 20, 10) from the first to the last layer with bagging size 100. For the fsDNN method, we set the cutoff p-value for important features to 0.1. Results are presented in Table 2, and the receiver operating characteristic curves (ROCs) for classifying acute POP status are displayed in Figure 1.

**Table 2:** Performance Comparison on Acute POP Data Analysis

| Method | Accuracy | AUC | MSPE<sub>1</sub> | MSPE<sub>2</sub> |
| --- | --- | --- | --- | --- |
| Parametric | 0.751 | 0.867 | 15.82 | 10.99 |
| Semiparametric | 0.751 | 0.867 | 15.88 | 10.84 |
| sDNN | 0.878 | 0.935 | 8.000 | 6.163 |
| fsDNN | 0.880 | 0.935 | 7.934 | 6.124 |

**Fig 1:**
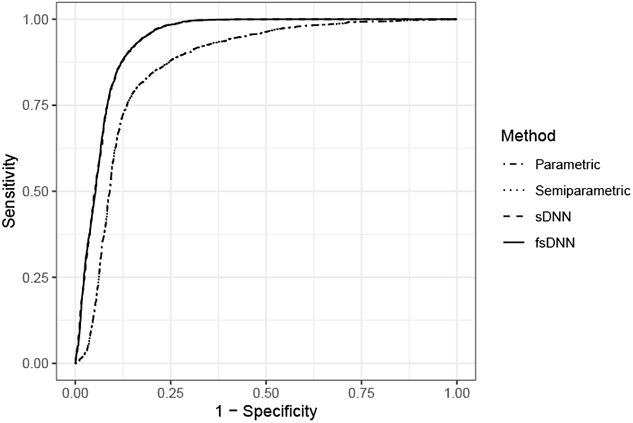
Classification Accuracy Comparison

The results from Table 2 and Figure 1 clearly demonstrate that the Parametric and Semiparametric methods are limited in their abilities to capture the complex data structures embedded in POP data, as they suffer from markedly worse performance in terms of their ability to classify acute POP status and to predict POP intensity compared to the proposed sDNN-based two-part predictive modeling procedures. For both the Paramateric and Semiparametric methods, the classification accuracy and AUC metrics were 0.751 and 0.867, respectively markedly inferior to the corresponding sDNN values of 0.878 and 0.935. The conditional and marginal mean squared prediction error (i.e., MSPE_1_ and MSPE_2_) based on the Parametric (Semiparametric) methods are almost twice and more than one and half times that of sDNN 15.82(15.88) vs 8.000 and 10.99(10.84) vs 6.163, respectively. It should also be noted that in addition to its inferior performance, the Semiparametric method using the kernel technique entails a much more intensive computational workload compared with the sDNN and fsDNN methods. Although in this dataset the improvement of fsDNN over sDNN is only marginal for all performance metrics considered, fsDNN offers robust statistical inference to assess the role of each feature associated with the acute POP outcomes. The clear dominance of our method recommends its utility in clinical practice for the purpose of POP management.

## 4. Simulation Studies

To explore the applicability and evaluate the performance of the proposed sDNN and fsDNN two-part predictive modeling framework for different types of semi-continuous data, we conducted numerical studies under two scenarios that captured a wide range of data structure complexity. For these scenarios, we allowed the feature matrix, ***X***, to be comprised of both continuous and discrete covariates, denoted as ***x*** and ***w***, repectively, such that each *x*_*p*_ ∼ *N* (0, 1), and each *w*_*p*_ ∼ *Bern*(0.5). A subset of each set of covariates of size 10 was assumed to affect disease status and severity in each scenario as follows:

### Scenario 1 (Truncated-normal data with simple structure)

In this scenario, we considered a simple data structure where the sets of covariates influence both the binary disease status *z* and the continuous positive outcome *y* via models that contain only linear terms such that:

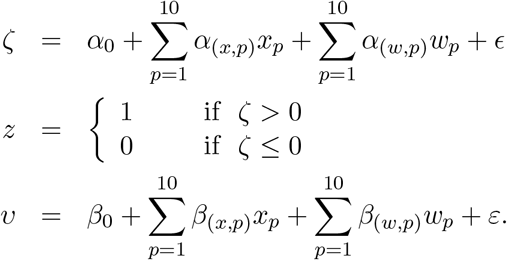

The positive continuous disease outcome *y* follows a truncated-normal distribution such that *y* = *υ* when *z* = 1 and *υ >* 0.

### Scenario 2 (Truncated-normal data with complex structure)

Building on Scenario 1, we increased the data complexity by adding the quadratic terms and interactions of the linear terms, as follows:

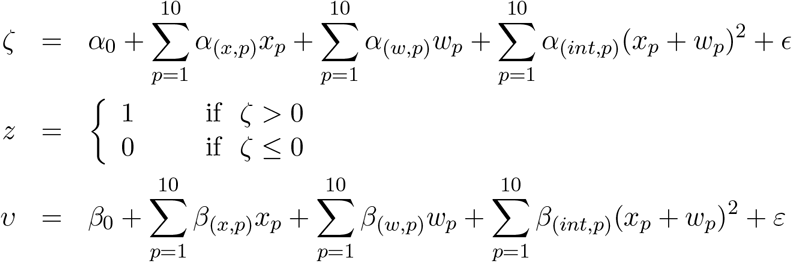

The positive continuous outcome *y* follows a truncated-normal distribution such that *y* = *υ* when *z* = 1 and *υ >* 0.

The random error terms, *E* and *ε* are assumed to be independent and to follow *N* (0, 1) distributions. Further, for each scenario, we ran two settings that differed in the number of nuisance covariates generated. In one scenario, we generated 15 additional continuous (∼ *N* (0, 1)) and 15 additional binary (∼ *Bern*(0.5)) nuisance covariates, which have no effect on the propensity of developing disease or the disease severity. In the second setting, we generated 40 of each such nuisance covariates. Thus, this resulted in either *p* = 50 or 100 observed covariates in total. The parameters *α*_0_, *α*_(*x*,1)_, · · ·, *α*_(*x*,10)_, *α*_(*w*,1)_, · · ·, *α*_(*w*,10)_, *α*_(*int*,1)_, · · ·, *α*_(*int*,10)_, *β*_0_, *β*_(*x*,1)_, · · ·, *β*_(*x*,10)_, *β*_(*w*,1)_, · · ·, *β*_(*w*,10)_, *β*_(*int*,1)_, · · ·, *β*_(*int*,10)_ are all drawn from standard normal distribution at the beginning of each simulation study and remain unchanged in the subsequent 1000 Monte Carlo replications.

Under the above simulation settings using sample sizes of both *n* = 1000 and 5000, we compare the proposed sDNN and fsDNN two-part predictive models to the Parametric and Semiparametric methods discussed in Section 3. In each Monte Carlo replication, we split the generated data into two parts such that 100 samples are reserved for the testing set and the rest are used for model training. For the sDNN and fsDNN methods, we use three hidden layers, with the number of hidden nodes (from the first to last layer) being 30, 20 and 10, with a bagging size of 100. Also, for the fsDNN method, the p-value cutoff is set to 0.1 to identify the important features for the two data processes to be included in the two predictive models.

The results for Scenario 1, with the number of observed covariates (*p*) set to 50 and 100, are summarized in Tables 3 and 4, respectively. While the performance of sDNN and fsDNN are nearly identical to the Parametric and Semiparametric methods for binary classification (regardless of sample size), they clearly dominate in terms of prediction accuracy for the continuous outcomes. For example, with 50 observed covariates and sample size 1000, the classification accuracy (AUC) from the Parametric, Semiparametric, sDNN, and fsDNN methods are 0.896(0.955), 0.896(0.955), 0.894(0.954), and 0.897(0.957), respectively. We note that the comparable binary classification performance of the DNN-based methods relative to the Parametric and Semiparametric methods is achieved despite the fact that these latter methods used multiple logistic regression models that are correctly specified in this simulation setting. The sDNN and fsDNN approaches both exhibit far superior performance in terms of prediction accuracy for the continuous outcomes (i.e., MSPE_1_ and MSPE_2_) relative to the other two approaches, regardless of sample size. The Parametric method is dominated because it is misspecified, employing multiple linear regression for the positive continuous outcomes, which were actually generated from truncated normal distributions. While the Semiparametric approach (which employs a single-index model with an unspecified link function approximated by the kernel method) is better than the Parametric approach for large sample size, it suffers from the curse of dimensionality - for the smaller sample size of 1000, it actually under-performs the Parametric method. Further, while both DNN-based approaches dominate in terms of prediction accuracy, fsDNN yields noticeable improvements in MSPE_1_ and MSPE_2_ above and beyond sDNN, resulting from its ability to effectively filter out the nuisance factors.

**Table 3:** Performance Comparison Under Scenario 1 (p = 50)

| Method | Accuracy | AUC | MSPE<sub>1</sub> | MSPE<sub>2</sub> |
| --- | --- | --- | --- | --- |
| Sample Size 1000 |  |  | Sample Size 1000 |  |
| Parametric | 0.896 | 0.955 | 8.414 | 3.771 |
| Semiparametric | 0.896 | 0.955 | 11.69 | 4.451 |
| sDNN | 0.894 | 0.954 | 6.075 | 3.327 |
| fsDNN | 0.897 | 0.957 | 5.811 | 3.195 |
| Sample Size 5000 |  |  | Sample Size 5000 |  |
| Parametric | 0.907 | 0.965 | 6.767 | 3.201 |
| Semiparametric | 0.907 | 0.965 | 5.429 | 2.936 |
| sDNN | 0.907 | 0.965 | 3.882 | 2.680 |
| fsDNN | 0.907 | 0.965 | 3.791 | 2.638 |

**Table 4:** Performance Comparison Under Scenario 1 (p = 100)

| Method | Accuracy | AUC | MSPE<sub>1</sub> | MSPE<sub>2</sub> |
| --- | --- | --- | --- | --- |
| Sample Size 1000 |  |  | Sample Size 1000 |  |
| Parametric | 0.878 | 0.939 | 11.09 | 4.906 |
| Semiparametric | 0.878 | 0.939 | 15.46 | 5.861 |
| sDNN | 0.887 | 0.948 | 6.881 | 3.571 |
| fsDNN | 0.894 | 0.954 | 6.347 | 3.323 |
| Sample Size 5000 |  |  | Sample Size 5000 |  |
| Parametric | 0.903 | 0.962 | 7.147 | 3.355 |
| Semiparametric | 0.903 | 0.962 | 6.963 | 3.320 |
| sDNN | 0.904 | 0.963 | 4.073 | 2.766 |
| fsDNN | 0.905 | 0.964 | 3.902 | 2.703 |

Analogous simulation results for Scenario 2 are presented in Table 5 (for *p* = 50) and Table 6 (for *p* = 100), respectively. Under this more complicated data structure that includes quadratic terms and interactions among the prognostic covariates, the dominance of the proposed sDNN and fsDNN methods applies not just to the prediction of continuous outcomes (as in Scenario 1), but extends to the classification of binary outcomes. Under these simulation settings, both the Parametric and Semiparametric methods are misspecified, leading to inaccurate classification for binary data and markedly imprecise prediction for continuous positive outcomes. For example, for sample size *n* = 5000 and observed covariate number *p* = 50, the classification accuracy (AUC) from the Parametric and Semiparametric methods are both 0.762 (0.809), compared to 0.909 (0.972) for sDNN. Even more striking, the MSPE_1_ (MSPE_2_) for the Parametric and Semiparametric approaches are 28.18 (28.55) and 21.40 (23.38), respectively, compared to 5.38 (8.17) for sDNN – reductions of 65-81% in favor of sDNN. Further, fsDNN yields substantial improvements relative to sDNN in terms of MSPE_1_(MSPE_2_) – on the order of 30% for sample size *n* = 1000 and number of covariates *p* = 50. As in Scenario 1, the curse of dimensionality hampers the performance of the Semiparametric method relative to the Parametric approach, despite the fact that the Semiparametric method relaxes the link function assumption; the Semiparametric approach only yields improved MSPE_1_ and MSPE_2_ for very large sample size.

**Table 5:** Performance Comparison Under Scenario 2 (p = 50)

| Method | Accuracy | AUC | MSPE<sub>1</sub> | MSPE<sub>2</sub> |
| --- | --- | --- | --- | --- |
| Sample Size 1000 |  |  | Sample Size 1000 |  |
| Parametric | 0.747 | 0.787 | 29.83 | 29.88 |
| Semiparametric | 0.747 | 0.787 | 37.14 | 32.42 |
| sDNN | 0.874 | 0.945 | 10.33 | 12.57 |
| fsDNN | 0.885 | 0.955 | 7.87 | 10.29 |
| Sample Size 5000 |  |  | Sample Size 5000 |  |
| Parametric | 0.762 | 0.809 | 28.18 | 28.55 |
| Semiparametric | 0.762 | 0.809 | 21.40 | 23.38 |
| sDNN | 0.909 | 0.972 | 5.38 | 8.17 |
| fsDNN | 0.910 | 0.972 | 5.21 | 7.91 |

**Table 6:** Performance Comparison Under Scenario 2 (p = 100)

| Method | Accuracy | AUC | MSPE<sub>1</sub> | MSPE<sub>2</sub> |
| --- | --- | --- | --- | --- |
| Sample Size 1000 |  |  | Sample Size 1000 |  |
| Parametric | 0.726 | 0.763 | 33.30 | 32.79 |
| Semiparametric | 0.726 | 0.763 | 46.91 | 38.99 |
| sDNN | 0.857 | 0.929 | 13.20 | 15.32 |
| fsDNN | 0.875 | 0.946 | 9.36 | 11.73 |
| Sample Size 5000 |  |  | Sample Size 5000 |  |
| Parametric | 0.758 | 0.804 | 28.36 | 28.63 |
| Semiparametric | 0.758 | 0.804 | 25.78 | 25.84 |
| sDNN | 0.905 | 0.968 | 5.64 | 8.45 |
| fsDNN | 0.909 | 0.972 | 5.25 | 7.91 |

To further investigate the robustness of the sDNN and fsDNN methods, we conducted two additional numerical studies with complex data structures which are modifications of Scenario 2 above, allowing for different types of positive outcomes *y*:

### Scenario 3 (Log-normal data with complex structure)

In this scenario, we consider a data structure where the sets of covariates influence both the binary disease status *z* and the continuous positive outcome *y* via models that contain a mixture of linear, quadratic, and interaction terms such that

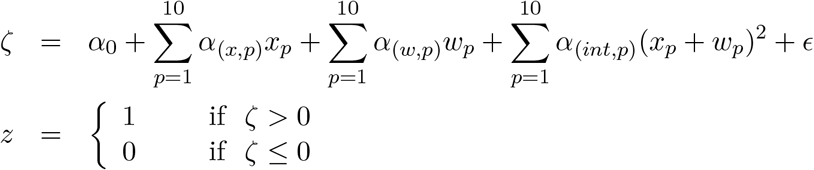

and the positive continuous disease outcome *y* is generated from the following log-normal model when *z* = 1:

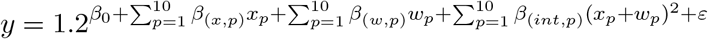

### Scenario 4 (Poisson data with complex structure)

Here, in addition to linear terms, we build on the models by adding the quadratic and interactions of the linear terms to influence the propensity of binary disease status *z*, as follows:

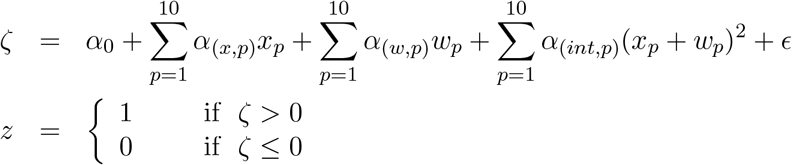

and the positive outcome *y* is generated from the following Poisson model with a mixture of linear, quadratic and interaction terms impacting the rate of the Poisson process when *z* = 1:

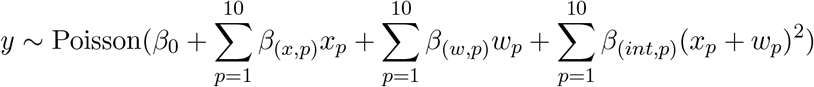

In these two scenarios, the important prognostic variables and nuisance features are generated as in Scenarios 1 and 2. The random error terms *E* and *ε* are assumed to be independent and to follow *N* (0, 1) distributions. The network structure used for the sDNN and fsDNN methods is the same as in Scenarios 1 and 2 as well. We ran 1000 Monte Carlo replications with sample size *n* = 1000 (of which 100 in each replication were randomly selected for testing) and observed number of covariates *p* = 50. These simulation results are displayed in Figure 2, and have been normalized by using the predicted MSPE_1_ and MSPE_2_ from the Semiparametric method to scale the other predicted MSPE_1_ and MSPE_2_ values.

**Fig 2:**
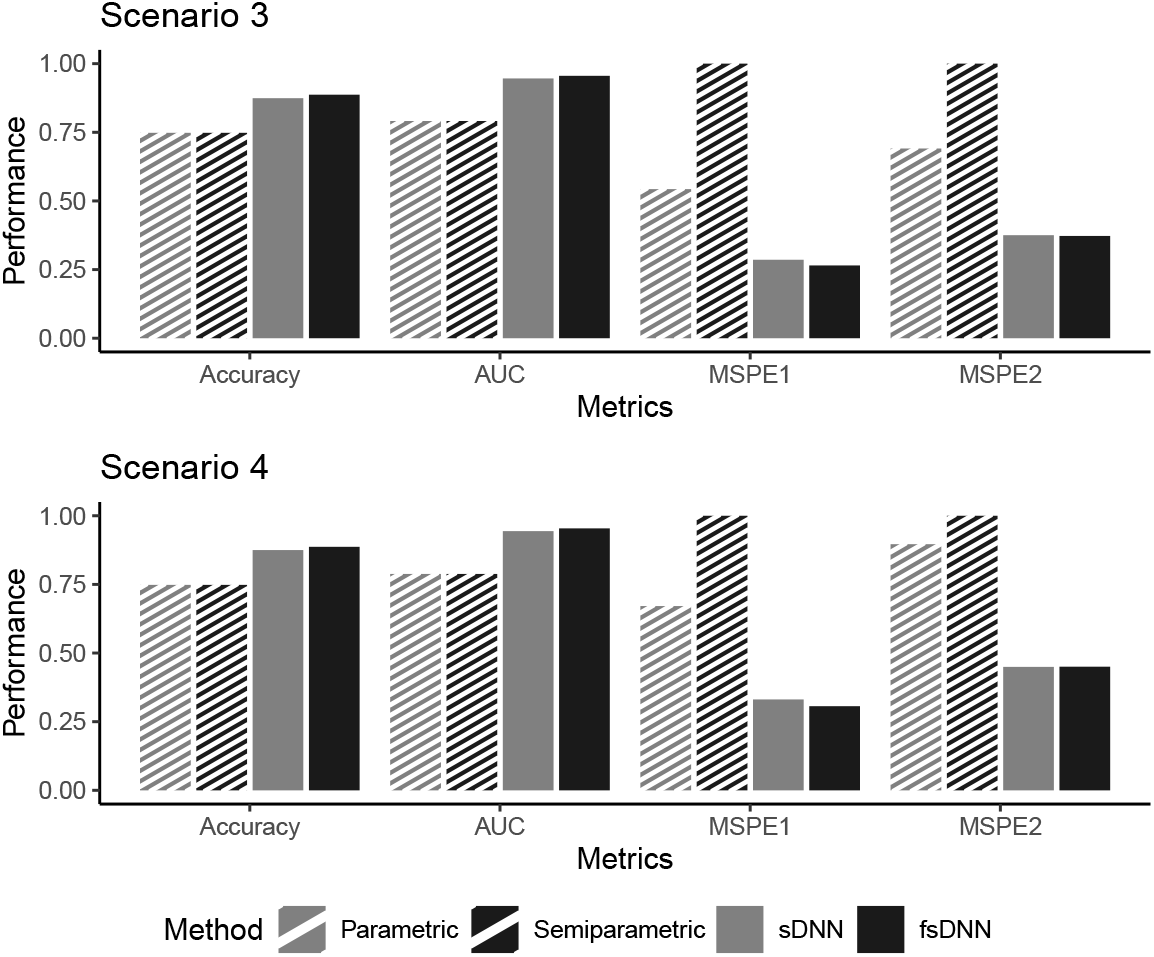
Performance Comparison for Scenarios 3 and 4

The results illustrated in Figure 2 are analogous to those for Scenario 2 (in Tables 5 and 6) in that the sDNN and fsDNN methods dominate the other two existing methods, achieving both markedly higher classification accuracy and substantially smaller prediction errors (for both metrics) regardless of whether the positive outcome is generated from a lognormal or Poisson model. Further, the fsDNN method also achieves some minor improvements over sDNN.

## 5. Discussion

In this paper, we derived a robust and powerful two-part predictive modeling procedure, sDNN, by adapting conventional DNN methods to model the complex architecture of postoperative pain data. We further expanded sDNN to fsDNN by incorporating a feature importance identification framework to provide a valid test for each included factor’s contribution to the accuracy of the semi-continuous outcome prediction. Although these proposed DNN-based two-part modeling methods were specifically motivated by the challenging POP data, our extensive numerical studies clearly demonstrate the advantages of the proposed methods relative to existing methods for a very general class of semi-continuous data that frequently appears in a range of scientific studies. Our methods are highly efficient for simple data structures that meet the assumptions of parametric models, and significantly more powerful for complex data structures for which the parametric assumptions fail. Our conclusions are robust to the number of variables modeled. These results clearly demonstrate the generality and effectiveness of DNN-based two-part modeling of semi-continuous data. It is worth pointing out that clinical outcome measurements can be correlated. For example, they may be collected longitudinally wherein correlation manifests within subjects. Alternatively, cross-sectional data can be correlated due to, for example, common environmental hazards or shared genetic structures among subjects. Robust yet efficient methods are needed for such correlated semi-continuous high-dimensional data, though this potential extension is beyond the scope of this paper.

## Notes

### Competing Interest Statement

The authors have declared no competing interest.

